## Supplementary figures and images for "Identification of Meibomian gland stem cell populations and mechanisms of aging"

### Fig. S1

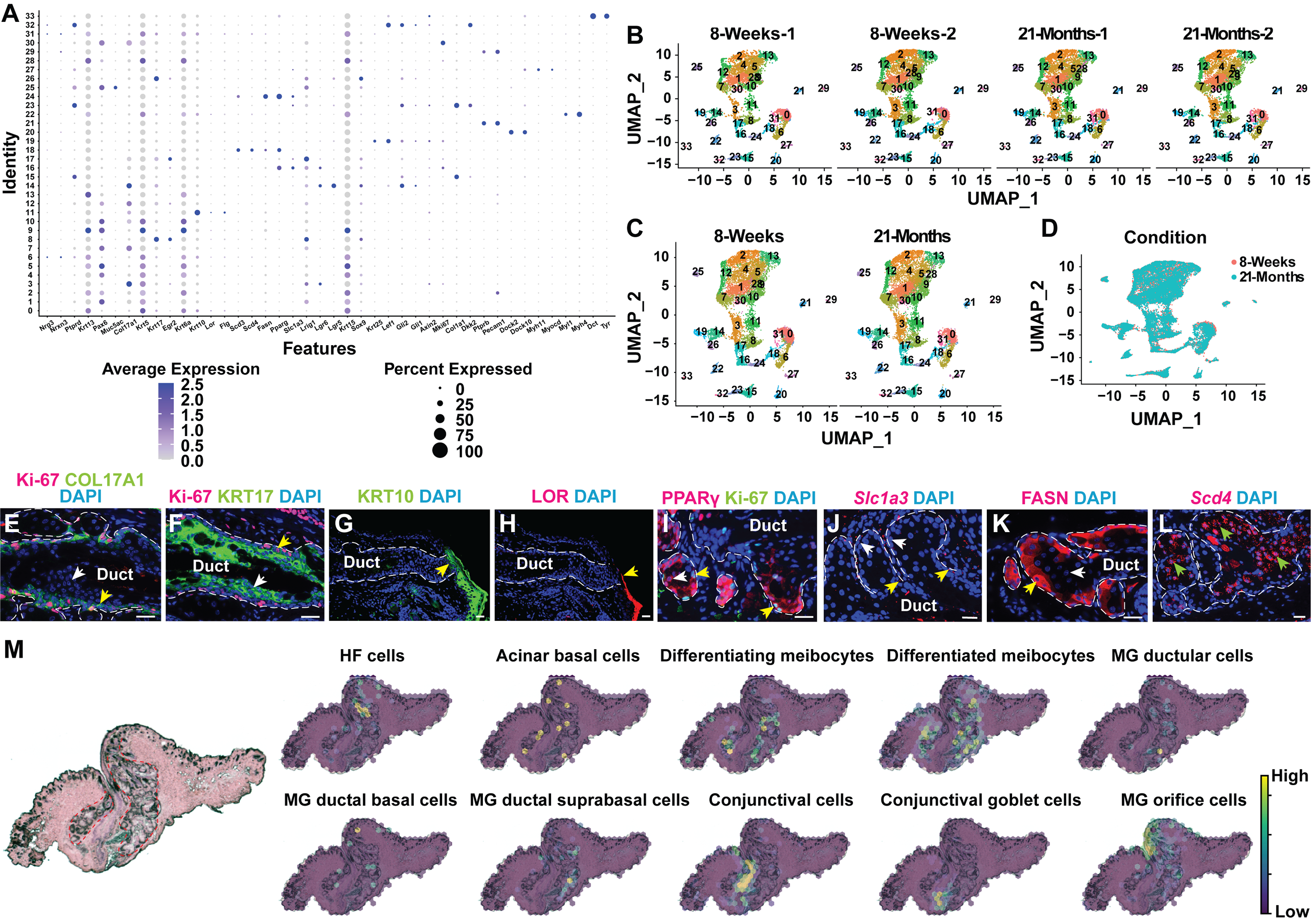

### Fig. S2

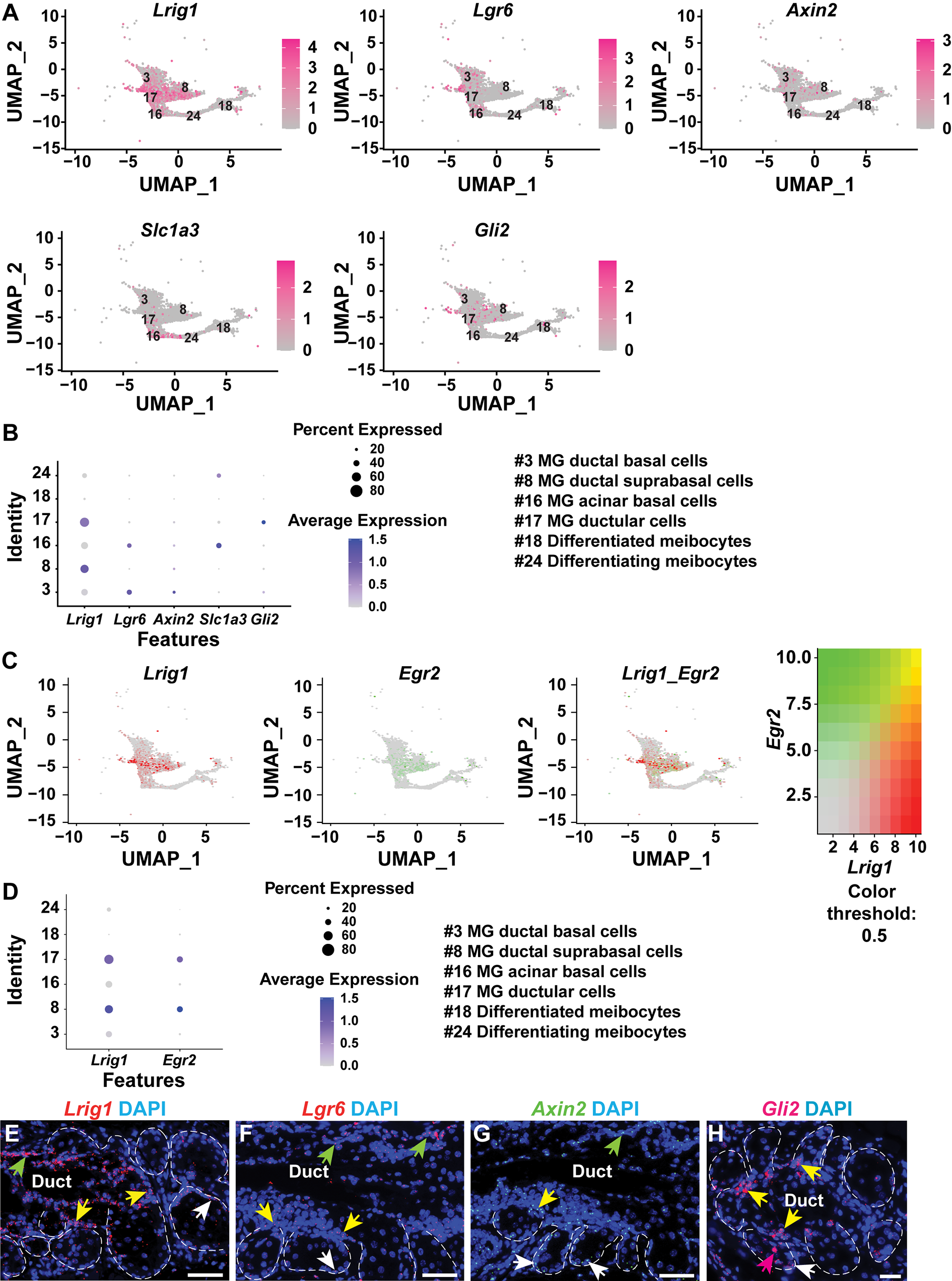

### Fig. S3

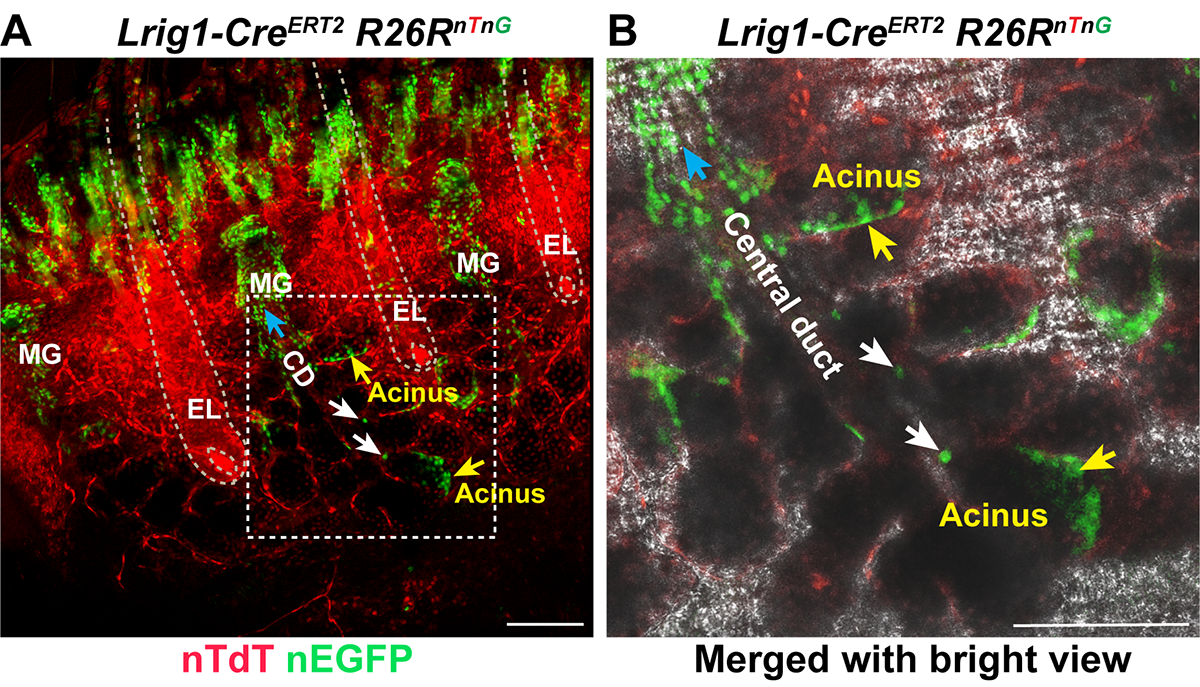

### Fig. S4

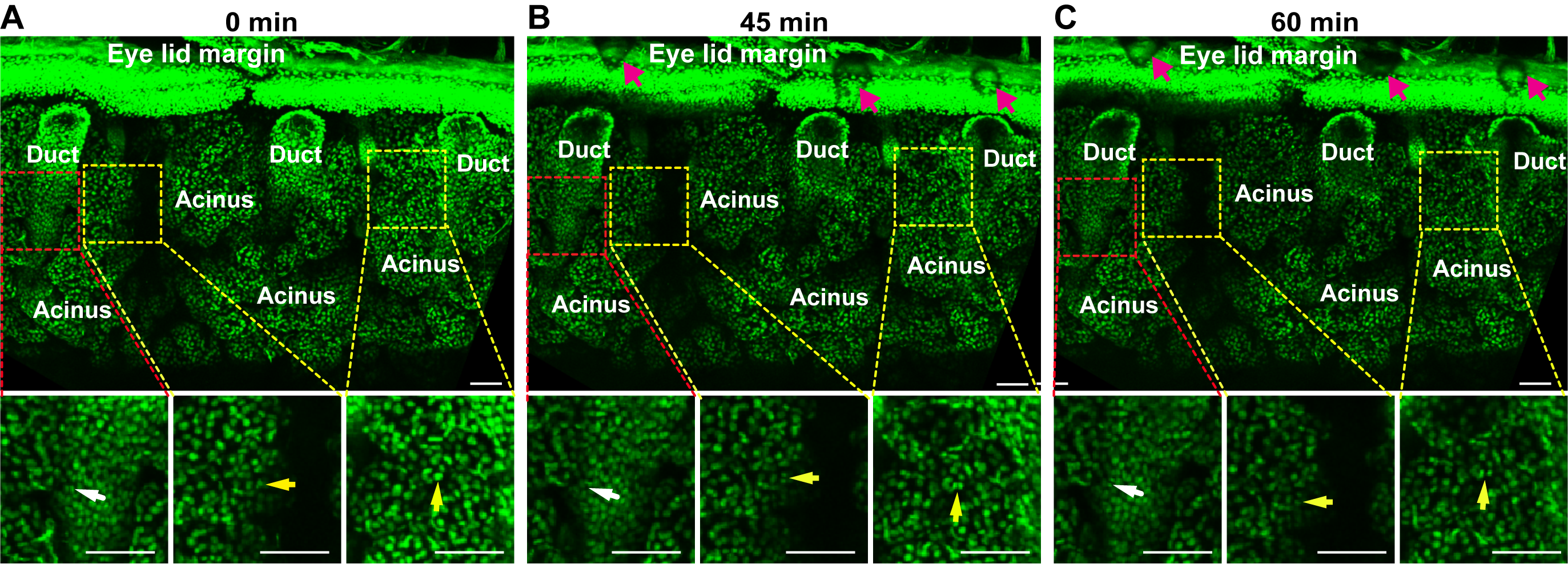

### Fig. S5

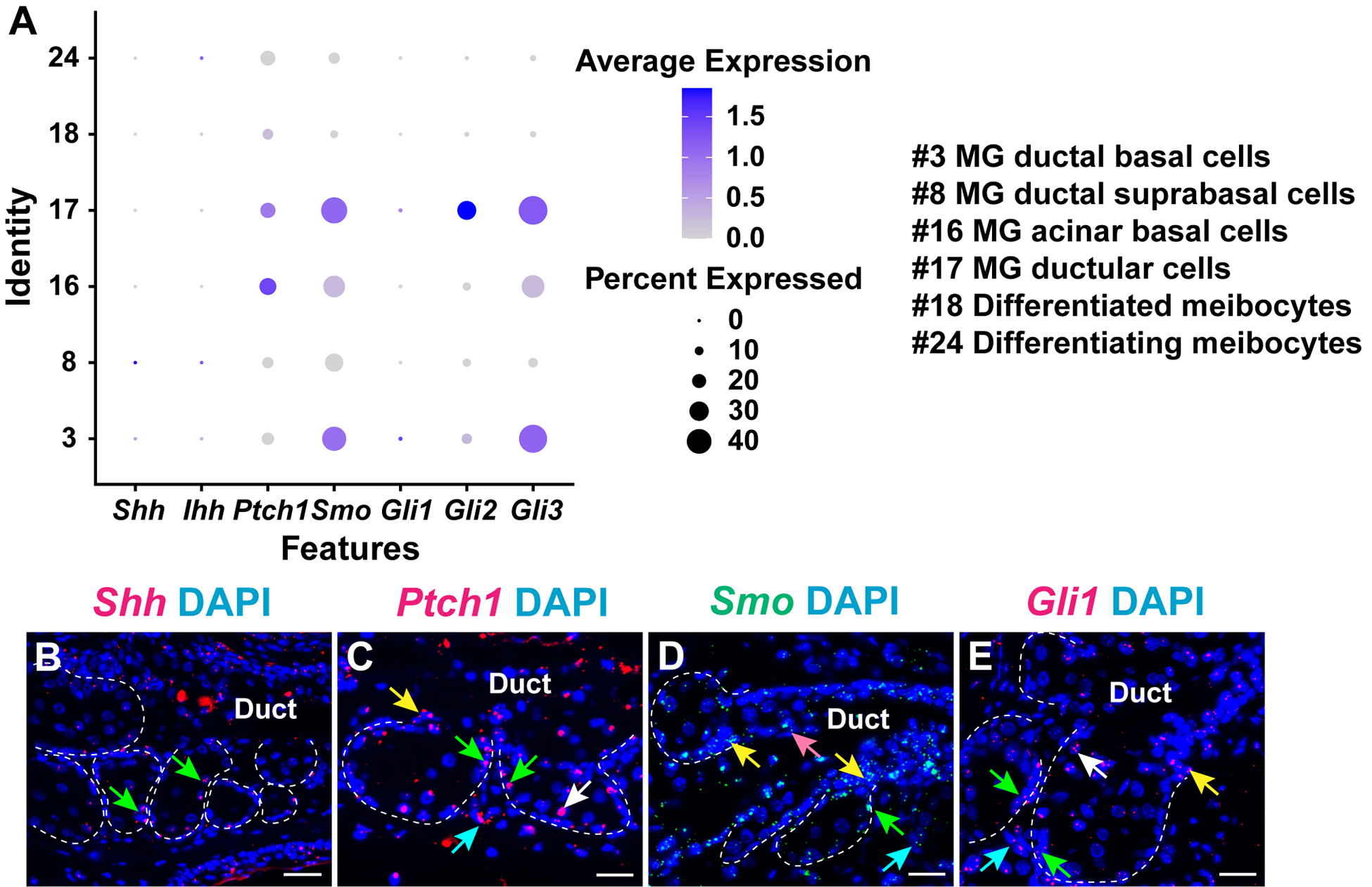

### Fig. S6

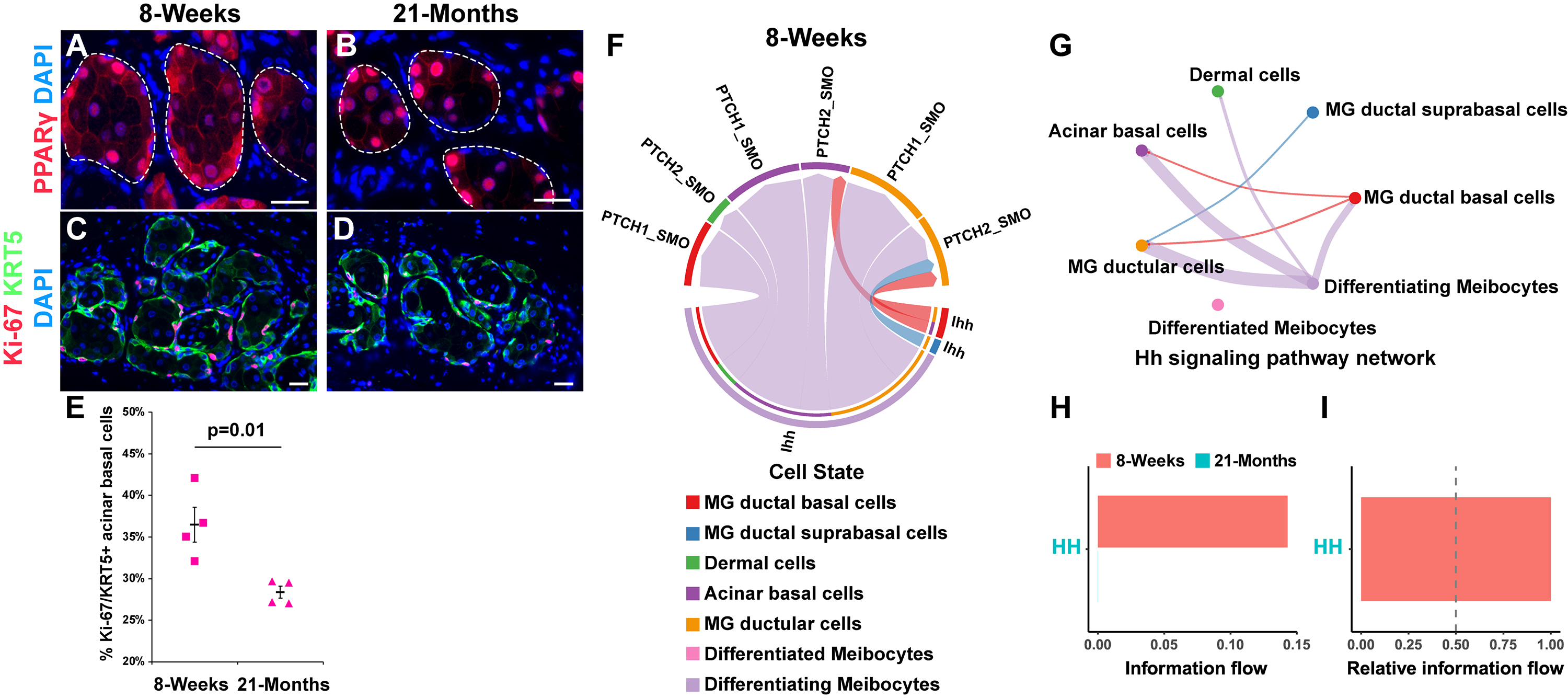

### Fig. S7

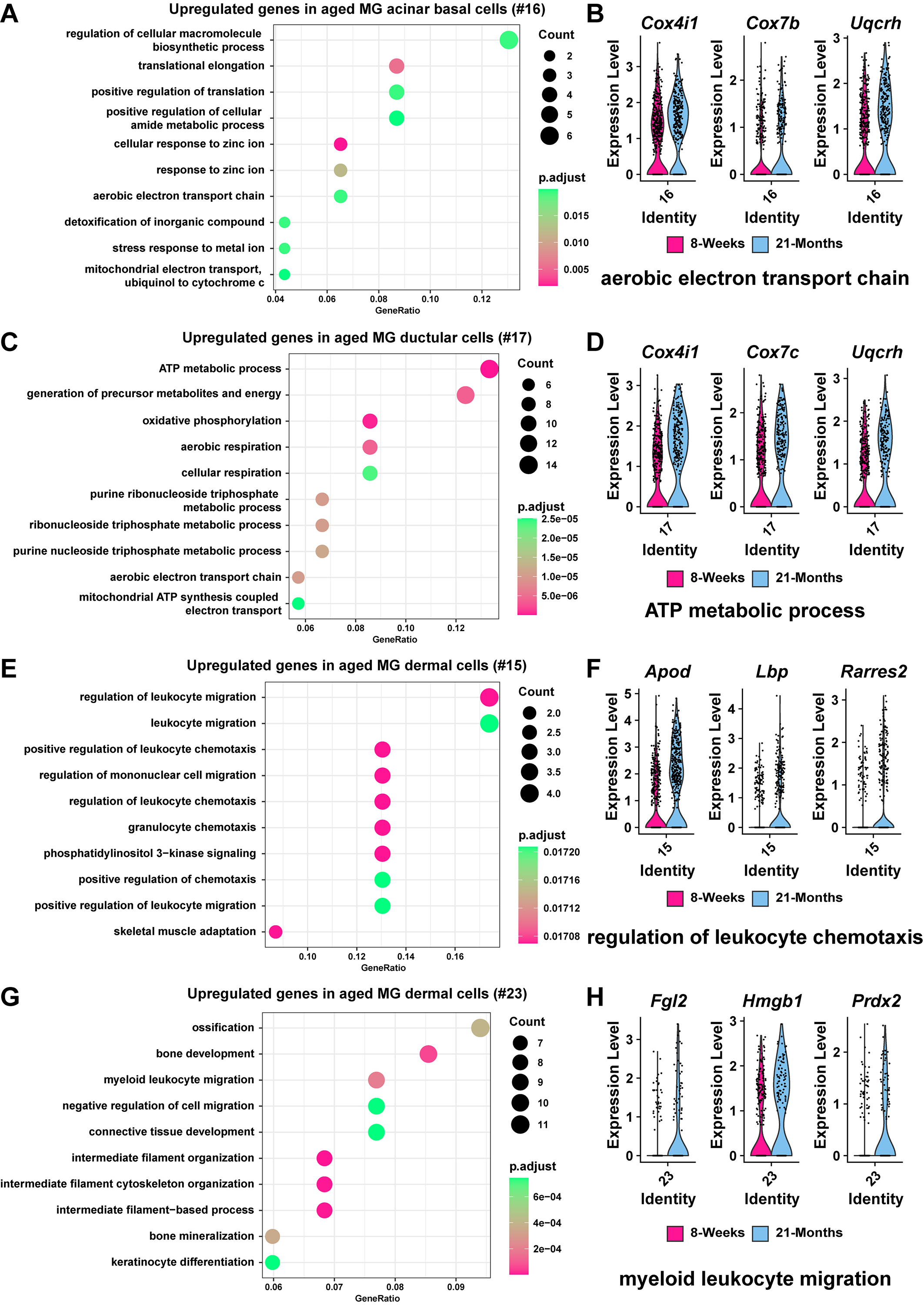
