## SUPPLEMENTARY FIGURE AND VIDEO LEGENDS for "Identification of Meibomian gland stem cell populations and mechanisms of aging"

**Supplemental Figure Legends**

**Figure S1. Identification of sub-populations of MG cells in snRNA-seq datasets and validation of cluster assignment via RNAscope and spatial transcriptomics.** (A) Dotplot shows the expression of signature genes for each cell cluster. (B) Separate UMAP plots for the four snRNA-seq datasets. (C) Separate UMAP plots for the combined replicate snRNA-seq datasets from 8-week-old and 21-month-old tarsal plates. (D) Merged UMAP plots of snRNA-seq datasets from 8-week-old and 21-month-old tarsal plates. (E) IF data show expression of COL17A1 in the Ki-67+ basal layer of MG ducts (yellow arrow) and its absence from the MG ductal suprabasal layer (white arrow). (F) IF for KRT17 shows that it is highly expressed in the MG ductal suprabasal layer (white arrow) and is weakly expressed in some ductal basal cells that are positive for Ki67 (yellow arrow). (G) IF shows that KRT10 is expressed in the ductal orifice (yellow arrow). (H) IF shows that LOR is expressed in the ductal orifice suprabasal layer (yellow arrow). (I) IF shows that PPARγ is expressed in Ki-67+ proliferative acinar basal cells (yellow arrows) and differentiating meibocytes (white arrow). (J) RNAscope for *Slc1a3* shows that it is primarily expressed in MG acinar basal cells (white arrows) and ductular cells (yellow arrows). (K) IF for FASN shows that it is expressed in differentiating meibocytes (yellow arrow) and not in fully differentiated meibocytes (white arrow). (L) RNAscope for *Scd4* shows that it is predominantly expressed in differentiated meibocytes (green arrows). White dashed lines in (E-L) outline the MG duct and/or acinus. Scale bars in (E-L) represent 25 μm. 3 independent samples were analyzed for IF and RNAscope. (M) Spatial transcriptomics analysis of 8-week MG using the 10x Visium platform; results were integrated with annotations based on the snRNA-seq data. Related to Figure 1.

**Figure S2. Expression of stem cell markers in the MG.** (A, B) Feature plots (A) and dot plot (B) from snRNA-seq data showing expression of *Lrig1*, *Lgr6*, *Axin2*, *Slc1a3*, and *Gli2* in MG cell populations. (C) Feature plots reveal partially overlapping expression of *Lrig1* and *Egr2* in MG cell populations. (D) Dot plot of snRNA-seq data shows enrichment of *Lrig1* and *Egr2* expression in MG duct suprabasal cells and ductular cells. (E) RNAscope validates that *Lrig1* is highly expressed in central duct (green arrow), moderately expressed in ductule (yellow arrows), and weakly expressed in the acinar basal layer. (F) RNAscope for *Lgr6* shows that it is broadly expressed in MG central duct (green arrows), ductule (yellow arrows), and acinar basal layer (white arrow). (G) RNAscope for *Axin2* shows that it is broadly expressed in central duct (green arrow), ductule (yellow arrow), and acinar basal layer (white arrows). (H) RNAscope for *Gli2* shows it is enriched in ductule (yellow arrows) and is also detected in the acinar basal layer (white arrow) and a sub-population of meibocytes (pink arrow). White dashed lines in (E-H) outline the MG acini. N=3 independent samples were used for RNAscope. Scale bars: (E-G), 50μm; (H), 25μm. Related to Figure 1.

**Figure S3. Progeny of *Lrig1+* stem cells are present in MG ductules, acini and central ducts.** (A) Whole mount confocal image of an MG explant from an *Lrig1-Cre^ERT2^ R26R^nTnG^* mouse induced at P36 and analyzed at P76. Lrig1 lineage traced cells (green) are present in ductules (white arrows), acini (yellow arrows) and central duct (blue arrow). (B) Enlarged view of the region outlined by a dashed white line in panel (A) and merged with bright view image. Grey dashed lines in (A) outline the eyelash follicles. N=2. EL, eyelash follicle; MG, Meibomian gland; CD, central duct. Scale bars: 150 μm. Related to Figure 1.

**Figure S4. MGs in tarsal plate explants undergo proliferation and secrete lipid droplets during short-term culture.** (A-C) Representative stills taken at 0 minutes (A), 45 minutes (B) and 60 minutes (C) from time-lapse live imaging of an adult K14:H2B-GFP^74^ murine MG explant in which all MG cells are GFP+. Areas shown by colored dashed boxes are shown at higher magnification beneath each image. Pink arrows indicate extruded drops of meibum. White arrows indicate a dividing cell in the central duct; yellow arrows indicate dividing cells in the acini. N=2. Scale bars: 100μm. Related to Figure 1.

**Figure S5. Hh pathway components are expressed in the MG.** (A) Dot plot of snRNA-seq data shows that Hh pathway components are expressed in MG subpopulations. (B-E) RNAscope (red) reveals expression of *Shh* in acinar basal cells (B, green arrows); *Ptch1* in acinar basal cells (C, green arrows), differentiating meibocytes (C, white arrow), ductular cells (C, yellow arrow), and stromal cells (C, light blue arrow); *Smo* expression in acinar basal cells (D, green arrow), ductular cells (D, yellow arrows), ductal basal cells (D, pink arrow), and stromal cells (D, light blue arrow); and *Gli1* in acinar basal cells (E, green arrows), differentiating meibocytes (E, white arrow), ductular cells (E, yellow arrow), and stromal cells (E, light blue arrow). White dashed lines in (B-E) outline acini. N=3. Scale bars: 25 μm. Related to Figure 2.

**Figure S6. Decreased proliferation and Hh signaling in aged MGs.** (A, B) IF data show that PPARγ is detected in the cytoplasm and nucleus of acinar cells in young MG (A) but is lost from the cytoplasm and enriched in the nucleus of acinar cells in aged MG (B). (C, D) Ki-67/KRT5+ proliferating acinar basal cells are reduced in aged (D) compared with young (C) MG. (E) Quantification shows that the percentage of KRT5+ basal cells that are Ki67+ is statistically significantly reduced in aged compared with young MGs (n=4 per condition). At least 114 acinar basal cells were analyzed per animal. Statistical significance was calculated using unpaired two-tailed Student’s t-test. Data are presented as mean +/- SEM. White dashed lines in (A, B) outline acini. (F, G) CellChat analysis indicating predicted Hh pathway-mediated intercellular communications among the indicated cell populations in young MG cells and between MG epithelial populations and surrounding dermal cells. Notably, no significant Hh pathway interactions were identified within aged MG cells or between aged MGs and surrounding dermal cells, due to extremely low Hh pathway gene expression in aged MGs. (H, I) CellChat analysis data depicting predicted intercellular crosstalk within MGs and between MGs and surrounding dermal cells mediated by Hh signaling pathways (H) and its relative strength in young and aged MGs (I). Related to Figure 5.

**Figure S7. GO analysis of upregulated genes in aged MG cells and surrounding dermal cells.** (A, B) GO analysis of genes with increased expression in aged compared with young acinar basal cells (cluster #16) (A) and violin plots for aerobic electron transport chain genes in acinar basal cells (B). (C, D) GO analysis of genes with increased expression in aged compared with young MG ductular cells (cluster #17) (C) and violin plots for oxidative phosphorylation pathway genes in ductular cells (D). (E, F) GO analysis of genes with increased expression in aged compared with young cluster #15 dermal cells (E) and violin plots for leukocyte chemotaxis genes in cluster #15 dermal cells (F). (G, H) GO analysis of genes with increased expression in aged compared with young cluster #23 dermal cells (G) and violin plots for myeloid leukocyte migration genes in cluster #23 dermal cells (H). Related to Figure 7.

**Video S1. MGs in tarsal plate explants undergo proliferation and secrete lipid droplets during short-term culture.** Time-lapse confocal live imaging of an adult K14:H2B-GFP murine MG explant in which all MG cells are GFP+. Note extrusion of droplets of meibum. Red arrow indicates a dividing cell in the central duct; yellow arrows indicate dividing cells in the acini. N=2. Related to Figure 1.

**Video S2. Lrig1+ MG ductule stem cell progeny can migrate toward acinus and locate in the acinus basal layer.** Time-lapse confocal imaging over 15 hours of culture of an MG explant from an *Lrig1-Cre^ERT2^ ROSA^nTnG^* mouse induced from P40-P42 and analyzed at P102. The white arrowhead labels an Lrig1+ stem cell migrating from the ductule towards the acinus. The red arrowhead marks a ductular Lrig1+ stem cell that divides once and then both daughter cells, indicated respectively with green and yellow arrowheads, migrate toward the acinus. The central duct is outlined with white dots; the acini are outlined with red dashed lines. Related to Figure 1.
